# Positive feedback promotes terrestrial emergence behaviour in an amphibious fish

**DOI:** 10.1101/2021.11.29.470419

**Authors:** Liam Tigert, Andy J. Turko, Patricia A. Wright

## Abstract

Major ecological transitions such as the invasion of land by aquatic vertebrates have been hypothesised to be facilitated by positive feedback between habitat choice and phenotypic plasticity. We tested whether aquatic hypoxia, emergence behaviour, and plastic changes in gill surface area could create such a positive feedback loop and drive an amphibious fish to spend increasing amounts of time out of water. We found that terrestrially acclimated amphibious mangrove rivulus *Kryptolebias marmoratus* were more sensitive to, and less tolerant of, aquatic hypoxia relative to water-acclimated fish, which are necessary trade-offs for positive feedback to occur. Next, we acclimated fish to normoxic or hypoxic water with the opportunity to emerge for 7d to test the predictions that fish in hypoxic conditions should regularly leave water, reduce gill surface area, and become less hypoxia tolerant. Consistent with these predictions, fish in severe hypoxia spent almost 50% of the time out of water and coverage of the gill lamellae by an inter-lamellar cell mass almost doubled. Hypoxia acclimated fish were also more sensitive to acute aquatic hypoxia (emergence at higher oxygen levels), and lost equilibrium faster in hypoxic water compared to control fish. Thus, we show that a positive feedback loop develops in amphibious fish where emergence behaviour begets further emergence behaviour, driven by gill remodelling which reduces aquatic respiratory function. Such a scenario may explain how amphibious behaviour has repeatedly evolved in fishes that occupy hypoxic aquatic habitats despite the associated challenges of life on land.

## Introduction

Spatial and temporal environmental heterogeneity creates a fundamental phenotype-environment mismatching problem for animals, as phenotypes well-suited to one environment are often poorly suited to different environments (Agrawal, 2020). To minimize these functional trade-offs, animals that inhabit variable environments either adopt generalist “jack-of-all-trades” phenotypes or are responsive to environmental conditions (Kassen, 2002; West-Eberhard, 2003). One form of responsiveness is phenotypic plasticity, the ability of an animal to modify its phenotype to match prevailing conditions (Pfennig, 2021). Another form of environmental responsiveness occurs when animals respond to variability by choosing to occupy the most favourable subset of available habitats (Edelaar et al., 2008). However there is little empirical data about how plasticity and habitat choice interact to determine phenotypes in heterogeneous environments (Edelaar and Bolnick, 2019).

Phenotypic plasticity and habitat choice can theoretically negate or increase the strength of the other process. For an organism that can always select an optimal habitat, there is no need for plasticity to improve the environment-phenotype match (Scheiner, 2016; Edelaar et al., 2017). Similarly, phenotypically plastic animals are often habitat generalists, as phenotypes can easily be adjusted to suit various conditions (Asbury and Adolph, 2007; Manenti et al., 2013). However, plasticity and habitat choice can also reinforce one another via positive feedback, although empirical examples are rare (Dingemanse and Wolf, 2010; Miner et al., 2005; West-Eberhard, 1989). In stream-living salamanders, individuals choose to inhabit fast-flowing riffles or slow-moving pools depending on morphological traits linked to swimming ability; phenotypic plasticity then reinforces these morphological differences between habitats (Lowe and Addis, 2019). Similarly, predatory ambush bugs choose to perch on flowers that are a similarly coloured to their own bodies, and then this match is improved via plastic changes to body colouration (Boyle and Start, 2020). These examples of habitat choice-plasticity feedbacks focus on spatially heterogeneous but temporally consistent environments. In many cases, however, environmental conditions vary in both space and time, but to date there has been little consideration of how plasticity and habitat choice interact in these scenarios.

In the face of temporal environmental change, animals must evaluate whether to remain and use plasticity to mitigate the phenotype-environment mismatch or maintain their morphological phenotype and move elsewhere (Hendry, 2016; Scheiner 2016). If environmental changes are gradual, plastic responses that produce a more “tolerant” phenotype may be expected given that dispersal often has fitness costs (Edelaar et al., 2008). However, for many environmental variables, there are probably thresholds of change that cannot be managed via plasticity and therefore require avoidance behaviour. In these situations, previous acclimation to the changing environment often results in the expression of a specialist phenotype which may be poorly suited to alternative environmental conditions due to functional trade-offs. Plastic expression of an alternative phenotype in the new environment is thus expected, and as the phenotype-environment match improves, a return to the previously occupied habitat should become less likely. In this way, the existence of habitat choice-plasticity feedback may reduce the occurrence of habitat transitions but also increase their success.

One of the most dramatic environmental transitions experienced by any animal occurs when amphibious fishes move between water and land (Ord and Cooke, 2016; Damsgaard et al., 2019; Turko et al., 2021). The fundamental physical differences between aquatic and terrestrial habitats result in a suite of functional trade-offs (Dejours, 1989). For example, fishes in water benefit from a large gill surface area that enhances oxygen uptake and improves tolerance to hypoxia, but in terrestrial conditions large gills may be a liability because of the potential for damage and evaporative water loss (Nilsson et al., 2012). This respiratory trade-off is especially problematic for amphibious fishes, as many species live in hypoxic aquatic habitats (Graham, 1997) and severe hypoxia is a common proximate reason why amphibious fishes leave water (e.g. Livingston et al., 2018; Mandic et al., 2009; Urbina et al., 2011; reviewed by Sayer, 2005). Once out of water, many amphibious fishes use alternative respiratory surfaces and gills become largely nonfunctional (Randall, et al., 1981). Some amphibious fishes, such as the mangrove rivulus *Kryptolebias marmoratus* (Ong et al., 2007) and bichir *Polypterus senegalus* (Turko et al., 2019a), modulate this gill surface area trade-off when moving between water and land by altering the size of the interlamellar cell mass (ILCM). In air, these fishes reversibly increase the ILCM, reducing overall gill surface area. However, upon the return to water, several days are required to decrease the ILCM and during this lag-time period respiratory function in water is impaired (Turko et al., 2012). Thus, by moving to land to escape aquatic hypoxia, and then developing a slowly reversed terrestrial gill morphology that limits oxygen uptake in water, a feedback loop may emerge that promotes terrestrial habitat choice (Turko et al., 2018).

We experimentally tested the hypothesis that habitat choice (behavioural hypoxia avoidance) and morphological plasticity (gill remodelling) reinforce one another in a positive feedback loop using the amphibious mangrove rivulus as a model. These fish readily move between water and land, survive prolonged episodes out of water, and experience severely hypoxic aquatic habitats in the wild (Taylor, 2012; Wright, 2012, Rossi et al., 2019). We first tested the critical predictions that terrestrial acclimation should decrease aquatic hypoxia tolerance and increase the sensitivity of the hypoxic emergence response. Next, we tested the prediction that hypoxia acclimation should cause emergence behaviour to become more frequent, and subsequent hypoxia tolerance should decrease (due to gill remodelling on land). In this hypoxia acclimation experiment, we observed that some fish remained in water despite extremely low oxygen saturation. To understand how these fish were able to obtain sufficient oxygen under these conditions, we quantified the usage of aquatic surface respiration (ASR) by mangrove rivulus across a range of dissolved oxygen concentrations.

## Methods

### Experimental Animals

Adult *K. marmoratus* hermaphrodites (0.09 g ± 0.02 g) originating from Belize (strain 50.91; Tatarenkov et al., 2010) and maintained in a breeding colony at the University of Guelph were used in these experiments. We kept fish individually in 120 ml clear plastic containers filled with 60 ml of brackish water (15‰, 25°C, 12-hour light:dark cycle), and fed them *Artemia* nauplii three times a week (Frick and Wright, 2002). Air exposure (7d) was achieved by placing fish on a brackish water-saturated filter paper substrate in standard housing containers (Ong et al., 2007). Control fish were held in identical containers filled with water and were fasted for the 7d acclimation period because mangrove rivulus are unable to feed while on land (Turko et al., 2019b; Wells et al., 2015).

### Experimental Protocol

#### Series 1 - Plasticity of hypoxia sensitivity and tolerance

To test how hypoxia sensitivity varied between air- and water-acclimated fish, we measured the acute emergence response of each acclimation group (water n=10, air n=10) as described previously (Regan et al., 2011, Livingston et al 2018). In brief, we placed individual fish into a two-part container, comprised of a lower section filled with water at 25°C, and an air-exposed platform that allowed fish to leave water. The fish were left in the lower section for 30 minutes to allow for recovery from handling stress, a time period determined to be sufficient in preliminary experiments. Atmospheric air was bubbled for the final 15 minutes of this acclimation period through an airline attached to the bottom of the water-filled container to acclimate the fish to gently bubbling water. We then remotely switched the air line to nitrogen gas and measured water oxygen saturation using a fibre optic oxygen probe (Witrox-4, Loligo Systems, Copenhagen, Denmark). We decreased the oxygen from 21 kPa to 0 kPa over 20 minutes and recorded the oxygen level at which the fish emerged.

To assess hypoxia tolerance, we measured time until loss of equilibrium (LOE) under severe hypoxia using standard methods (Regan et al., 2017). Briefly, we placed fish (water n=6, air n=6) into small, mesh sided chambers, and left them submerged in brackish water for 15 minutes to recover from handling stress. These chambers were then gently submerged into a 5-gallon glass aquarium to prevent the fish from reaching the air-water interface. The aquarium was covered with plastic film to prevent gas exchange with atmospheric air. Again, the fish were left to recover from handling stress for another 15 minutes, during which time oxygen concentration did not appreciably decrease. Then, we bubbled the water with compressed nitrogen gas to decrease the oxygen partial pressure of the water from 21 kPa to 0.5 kPa over 10 minutes, at which point it was held constant at 0.5 kPa for the remainder of the experiment (Regan and Richards, 2017; Regan et al., 2017). We recorded the time until the fish lost equilibrium, that is when the fish did not respond to the physical stimulus of three gentle prods.

#### Series 2 - Emergence behaviour and hypoxia sensitivity

To test the prediction that hypoxia acclimation should increase the frequency of emergence and subsequent decrease hypoxia tolerance, we acclimated fish (7d) to one of three oxygen levels: normoxia (21 kPa oxygen; n = 20), hypoxia (2.1 kPa oxygen, n =20) or extreme hypoxia (0.5 kPa; n = 15). Within each group, fish were acclimated to either water or air for 7d immediately before the experiment began to determine if differences in gill morphology caused by air-versus water-acclimation influence emergence behaviour in hypoxia. Fish were held individually in a two-part container with a platform for emergence as described previously (Livingston et al., 2018). These containers were similar to the ones used to assess the hypoxic emergence response (described above), except the lower, water-filled portion of the containers were mesh-sided and submerged in a larger water bath, allowing for free circulation of water maintained at one of three oxygen levels: We maintained oxygen levels within the basin with an oxygen control system (OxyCTRL, Loligo Systems, Copenhagen, Denmark) that automatically bubbled compressed nitrogen gas into a header tank when oxygen concentrations exceeded the experimental setpoint. We video recorded fish behaviour from above using a webcam during the 12 hours of light per day, as we previously found that there is no difference in emergence behaviour between day and night (Turko et al., 2011), and subsequently measured the proportion of time each fish spent out of water per day. At the conclusion of the 7d experiment, we tested the acute emergence response or time to LOE on random subsets of fish as described above.

We assessed gill surface area in a subsample of fish to determine if there was a relationship between behaviour and gill morphology. Fish were euthanized using MS-222 and processed for histology as described previously (Turko et al., 2011). We measured the height of the ILCMs in 5 randomly selected gill filaments for each fish and calculated the average proportion of ILCM cover per fish.

#### Series 3 - Aquatic surface respiration

We noticed that some fish in the extreme hypoxia treatment (0.5 kPa O2) never lost equilibrium despite remaining in water for lengths of time that far exceeded the degree of tolerance we measured in our LOE experiments, and hypothesised the fish were obtaining supplemental oxygen using aquatic surface respiration (ASR; Kramer and McClure, 1982). To test this hypothesis, we placed fish (air-acclimated n = 4, water-acclimated n=5) in a 30 cm upright mesh tube, inside of a 5-gallon glass aquarium filled halfway with water at 25°C and allowed them to recover for 15 minutes. The sides of the aquarium were covered, except for one viewing window, to limit external influences on fish behaviour. After the recovery from handling stress, we slowly introduced nitrogen gas into the water to reduce oxygen levels at a constant rate (−0.4 kPa oxygen/min), mixed thoroughly with stir bars, over the span of 50 minutes. Each trial was video recorded for its duration, and we subsequently quantified the proportion of time fish used ASR as a function of oxygen saturation in the water.

### Statistical Analysis

All statistics were calculated using R version 4.0.2. To compare oxygen saturation at emergence and time to LOE in water-versus air-acclimated fish, we used Student’s t-tests (Series 1). To compare oxygen saturation at emergence, time to LOE, and gill ILCM coverage in fish after acclimation to normoxic, hypoxic, or extremely hypoxic water (Series 2), we used two-way ANOVA, followed by Tukey’s post-hoc tests. To assess changes in emergence behaviour over time within each acclimation condition, we used permutational two-way ANOVA (1,000,000 permutations, package *permuco*; Frossard and Renaud 2019) because our data did not meet assumptions of normality or homoscedasticity. Permutational ANOVA is similar to a parametric ANOVA, but rather than assuming a Gaussian distribution, the dataset is repeatedly randomized, and the proportion of randomly ordered datasets with greater treatment effects than the actual dataset is calculated. We used a series of two-way ANOVAs (one per oxygen tension) rather than a single three-way ANOVA because we found a marginal 3-way interaction in our exploratory analysis (F = 1.73, p = 0.056) and because the key predictions made by the positive feedback hypothesis relate to patterns over time within each treatment rather than statistical comparison among treatments. Post-hoc comparisons were calculated using permutational t-tests (1,000,000 permutations) implemented using the pairwise.perm.t.test function (package *RVAideMemoire,* Hervé 2020), and p-values were adjusted for multiple comparisons using the Benjamini and Yekutieli correction which controls the false discovery rate (Benjamini & Yekutieli, 2001). We used similar permutational ANOVA and post-hoc permutation t-tests (with Benjamini and Yekutieli corrected p-values) to assess changes in the proportion of time fish used aquatic surface respiration during the induction of hypoxia (Series 3), as these data did not meet parametric assumptions of normality or homoscedasticity.

## Results

The hypoxic emergence response of air-acclimated fish was almost twice as sensitive as that of water-acclimated fish (t-test p = 0.004; Fig. 1A). Hypoxia tolerance (time to LOE) of air-acclimated fish was approximately half that of water-acclimated fish (t-test p = 0.0007; Fig. 1B).

**Figure 1.**
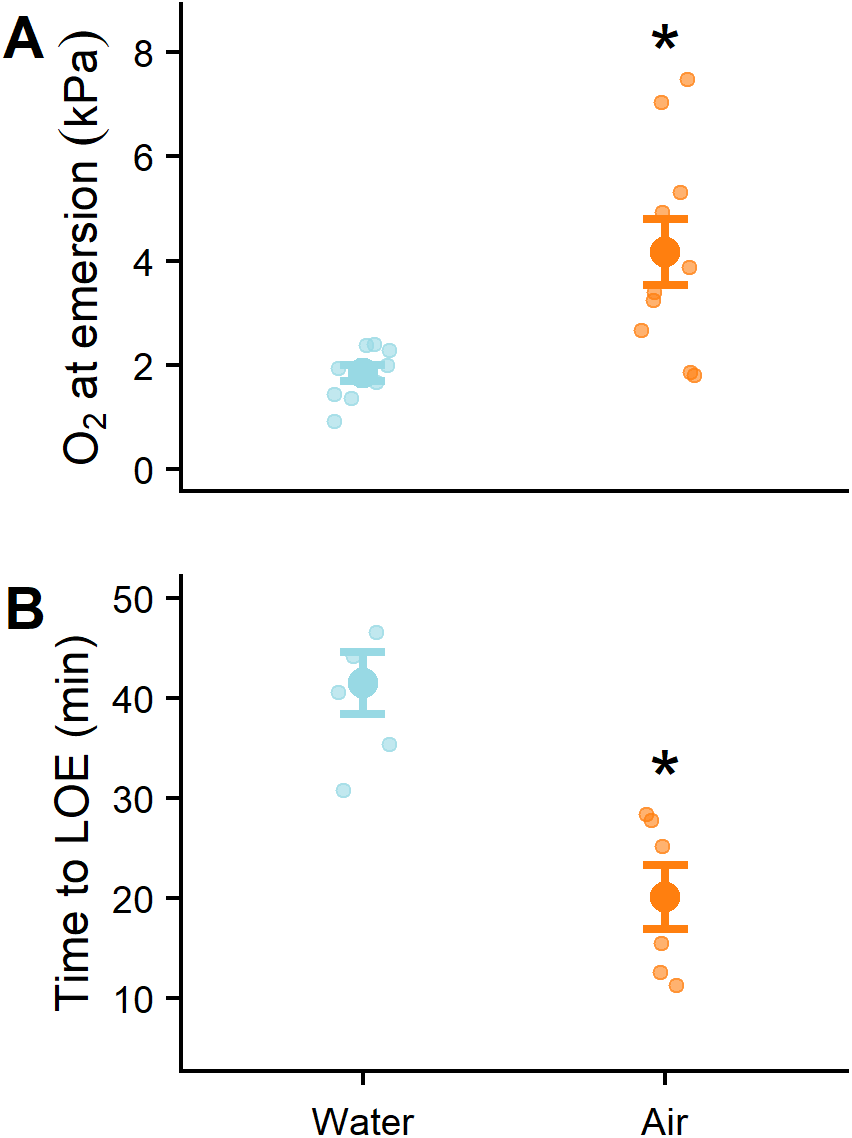
Sensitivity and tolerance of *Kryptolebias marmoratus* to aquatic hypoxia after acclimation to water or air. (A) Partial pressure of oxygen at the point of emergence (n=10/treatment). (B) Time until loss of equilibrium at 0.5 kPa (n=6/treatment). Water-acclimated fish are shown in blue, air-acclimated fish are shown in orange. Small points denote raw values, large points denote group means, and error bars show standard error. An asterisk denotes significant difference between acclimation treatments (p<0.05).

In our habitat choice experiment, exposure to aquatic hypoxia dramatically increased the amount of time fish spent out of water (Fig. 2). Fish in normoxia rarely emerged from water over the course of the trial, and emergence frequency in previously air-acclimated fish slightly but significantly decreased over the 7d trial (time * acclimation p = 0.0069; Fig. 2A). In moderate hypoxia, emergence frequency similarly decreased over time, and previously air-acclimated fish initially spent significantly more time out of water than water-acclimated fish (time * acclimation p = 0.0072; Fig. 2B). In extreme hypoxia, fish regularly emerged from water (~30% of the time) but this did not depend on previous acclimation condition (p = 0.12) nor time (p = 0.057, interaction p = 0.11; Fig. 2C).

**Figure 2.**
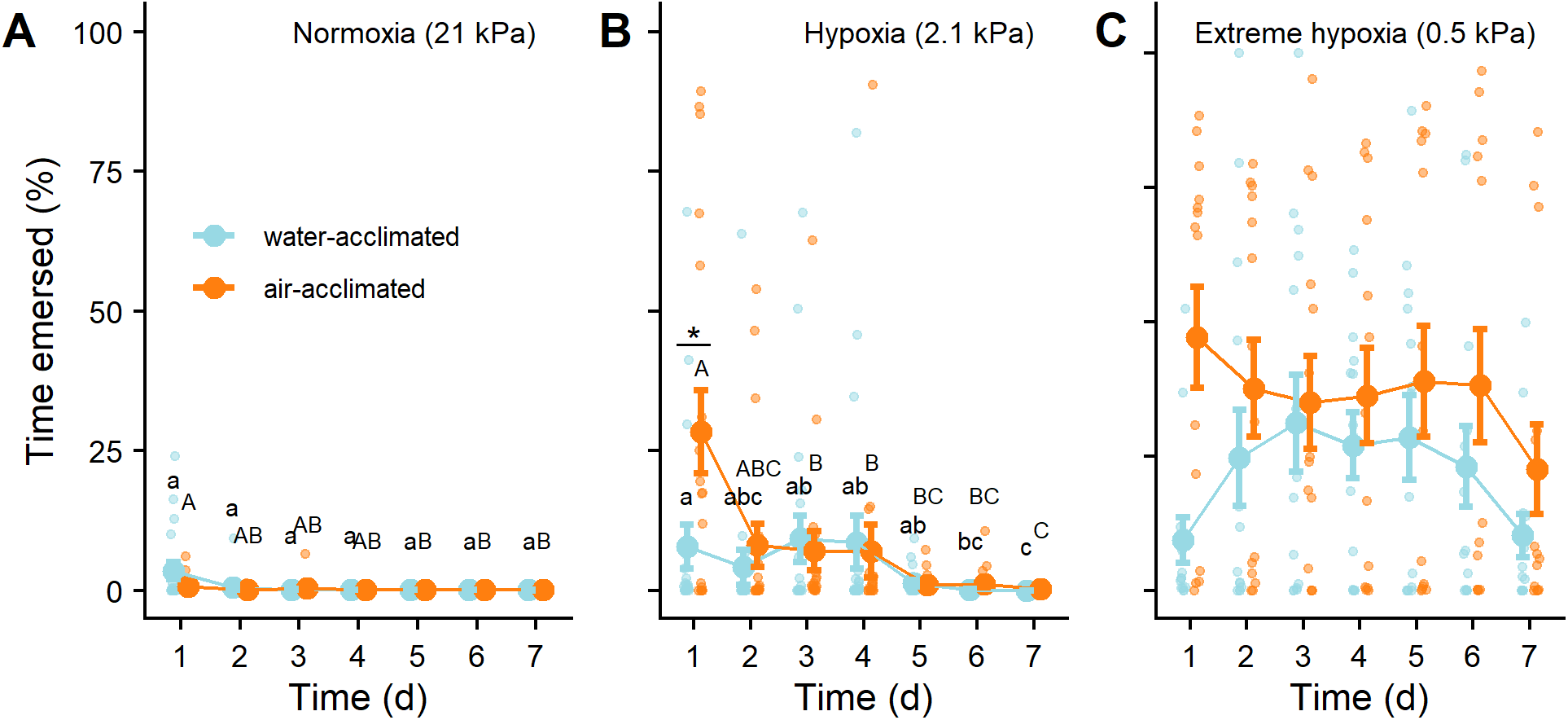
Habitat choice in *Kryptolebias marmoratus* acclimated to varying levels of aquatic hypoxia. Proportion of time air- or water-acclimated fish spent on land during exposure to water of one of three oxygen pressures: (A) normoxia (21 kPa; n=20), (B) hypoxia (2.1 kPa, n=20, and (C) extreme hypoxia (0.5 kPa, n=15). Water-acclimated fish are shown in blue, air-acclimated fish are shown in orange. Small points denote raw values, large points denote group means, and error bars show standard error. Different letters within an acclimation group denote statistical differences over time (p < 0.05). An asterisk denotes significant difference between acclimation treatments (p<0.05).

At the end of the habitat choice experiment, hypoxia-acclimated fish had a significantly more sensitive hypoxic emergence response compared to fish acclimated to normoxic conditions (two-way ANOVA, p < 0.0001; Fig. 3A). Acclimation to aquatic hypoxia also decreased hypoxia tolerance (time to LOE) by about 50% (two-way ANOVA, p < 0.00001; Fig. 3B). Finally, acclimation to extreme aquatic hypoxia almost doubled the proportion of gill lamellae that were covered by an inter-lamellar cell mass compared to normoxic conditions (two-way ANOVA, p = 0.002; Fig. 2C). In all cases, there was no significant effect of previous acclimation to air versus water (all p > 0.05).

**Figure 3.**
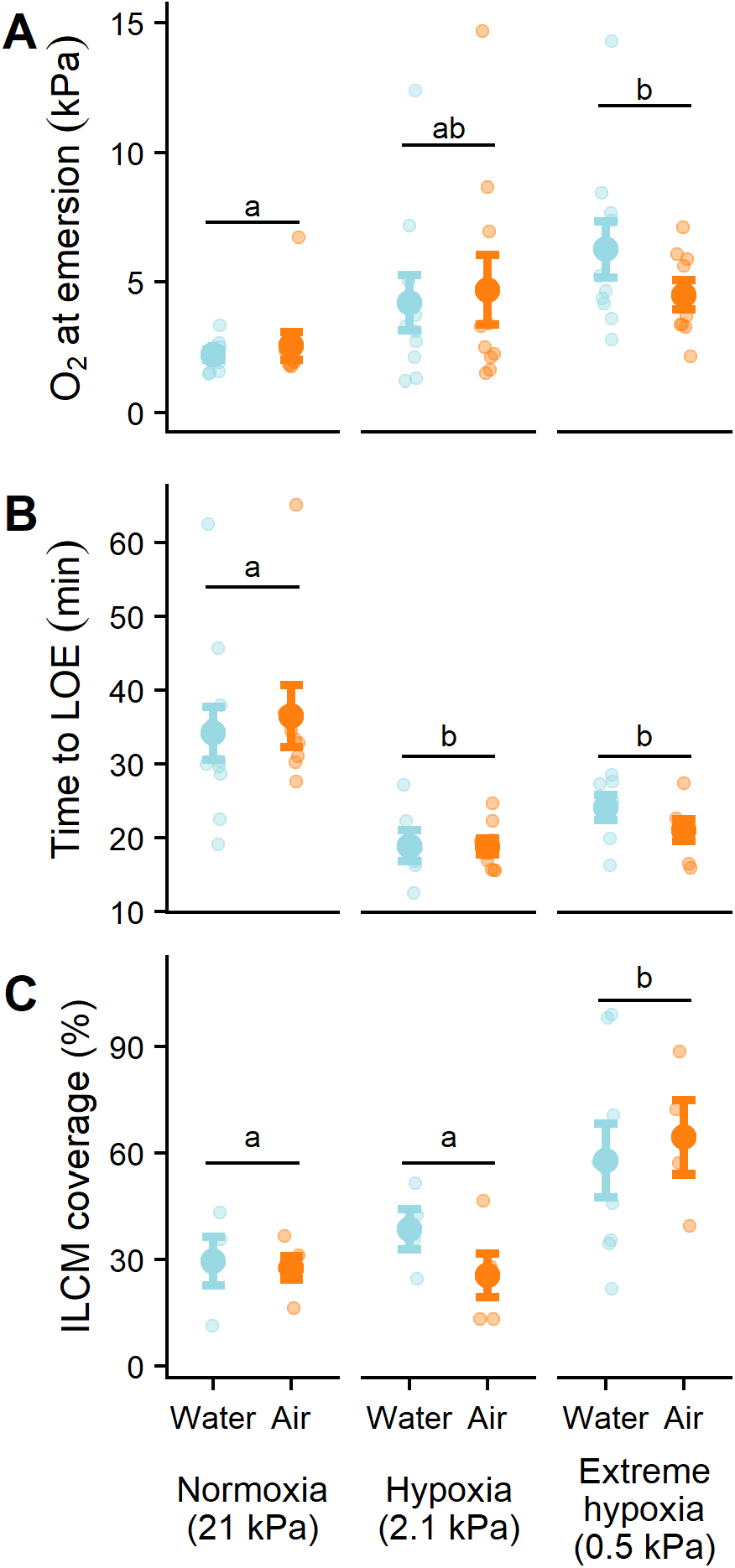
Phenotypic plasticity of *Kryptolebias marmoratus* after acclimation to varying levels of aquatic hypoxia with access to terrestrial habitats. Water oxygen pressure at the point of emergence (A) and time until loss of equilibrium (B) and the proportion of the gill covered by an ILCM (C) between *K. marmoratus* acclimated to air or water for seven days, followed by a return to water with one of three oxygen pressures, over another seven days. Water-acclimated fish are shown in blue, air-acclimated fish are shown in orange. Small points denote raw values, large points denote group means, and error bars show standard error. Different letters denote a significant difference between oxygen pressure treatment.

Mangrove rivulus significantly increased their use of aquatic surface respiration when oxygen tension fell below 10 kPa (P<0.0005; Fig. 4). At the lowest P_O2_ we tested (1.5 kPa), fish used aquatic surface respiration for 95% of the recording period. This pattern of aquatic surface respiration was not affected by prior acclimation to air or water (P>0.05; Fig. 4).

**Figure 4.**
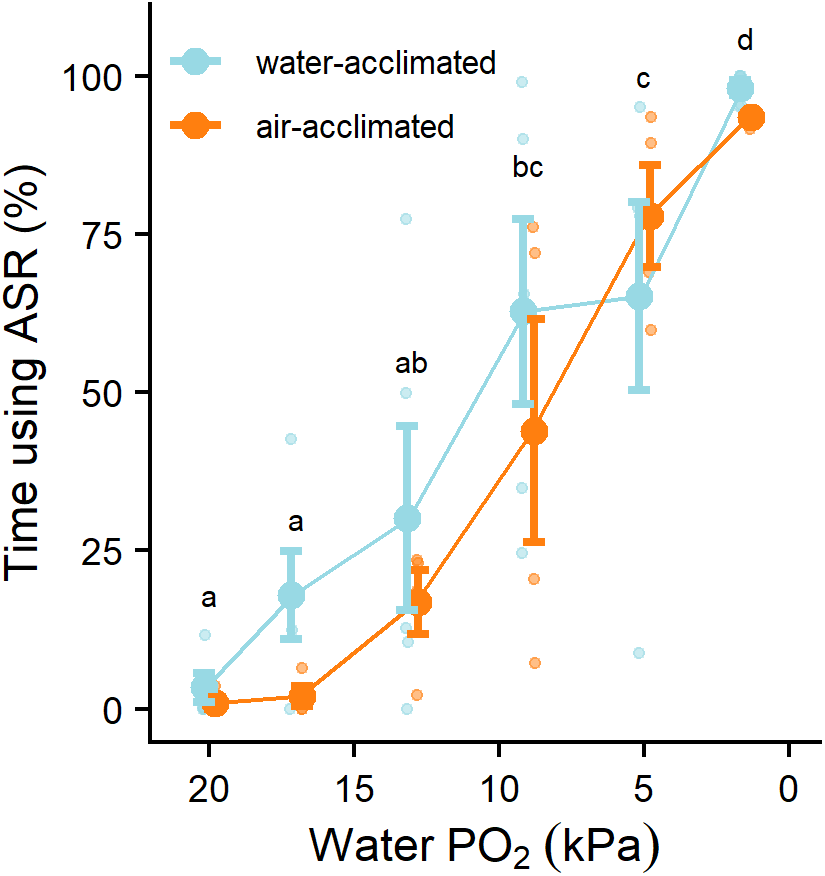
Use of aquatic surface respiration by *Kryptolebias marmoratus*. Proportion of time water- or air-acclimated fish spent at the surface in response to decreasing oxygen concentrations (n=6/treatment; note the reversed x-axis). Water-acclimated fish are shown in blue, air-acclimated fish are shown in orange. Small points denote raw values, large points denote group means, and error bars show standard error. Different letters denote overall statistical differences between levels of oxygen (p < 0.05).

## Discussion

Our results are generally consistent with the hypothesis that hypoxia-induced emergence behaviour becomes self-reinforcing via positive feedback as gill remodelling on land decreases respiratory function in water and promotes further emergence behaviour. We found three lines of evidence that support this hypothesis. First, air-acclimation increased the sensitivity to hypoxia as fish emerged at higher oxygen levels relative to water-acclimated fish. Second, when reintroduced to an aquatic environment, air-acclimated fish emerged more frequently and for longer durations than water-acclimated fish across all oxygen treatment groups. These are both necessary conditions for the establishment of a positive feedback loop. Finally, after a week of exposure to hypoxic water with access to land, hypoxia sensitivity increased while hypoxia tolerance decreased. Overall, our data indicate that emergence behaviour is an important strategy for coping with adverse aquatic conditions in this species, consistent with previous studies (Rossi et al., 2019; Turko et al., 2018). Importantly, our data also indicates that this emergence behaviour makes it increasingly difficult for these fish to maintain homeostasis in hypoxic water. We suggest that this sort of positive feedback between plasticity and habitat choice has widespread potential to generate extreme phenotypes in animals. Furthermore, we speculate that if habitat choice-plasticity feedback results in genetic assimilation (Crispo, 2007; Pigliucci et al., 2006; Schneider and Meyer, 2017), this process may accelerate evolutionary habitat transitions such as the invasion of land by fishes.

### Requirements for positive feedback

There are two key elements necessary to generate positive feedback between habitat choice and phenotypic plasticity. First, expressions of plasticity that improve performance in one habitat must also decrease performance in alternative habitats (i.e. an antagonistic trade-off). Second, an animal must be capable of assessing its own phenotype and using this information to choose suitable habitats (i.e. matching habitat choice; Camacho et al., 2020). Our data indicate that both elements exist for mangrove rivulus moving between water and land, suggesting that the positive feedback hypothesis is plausible.

Mangrove rivulus face a suite of physiological trade-offs when moving between aquatic and terrestrial environments, the first requirement for positive feedback to occur. Many of these trade-offs involve the oxygen transport cascade. Gill remodelling on land may reduce evaporative water loss in air but also decreases the capacity for oxygen uptake in water (Turko et al., 2012). Angiogenesis of epithelial capillaries on land enhances aerial oxygen uptake (Blanchard et al., 2019) perhaps at the cost of oxygen loss to hypoxic water (Scott et al., 2017). The oxygen binding affinity of hemoglobin also increases in air, but in water this may impair oxygen offloading to metabolically active tissues (Tunnah et al., 2021; Turko et al., 2014). Terrestrially induced hypertrophy of oxidative trunk muscle (Brunt et al., 2016; Rossi et al., 2018) may also detrimentally increase metabolic oxygen demand in hypoxic aquatic environments. In this study, we found that terrestrial acclimation decreased overall aquatic hypoxia tolerance (i.e. time to loss of equilibrium) by over 50%, indicating that the combined effects of these physiological trade-offs have a dramatic organism-level consequence.

We also found evidence that habitat choice depends on physiological state (i.e. self-assessment), fulfilling the second requirement for positive feedback. When acutely challenged with progressive aquatic hypoxia, air-acclimated fish emerged at oxygen concentrations that were more than double that of water-acclimated fish (1.8 vs. 4.2 kPa). This result implies that mangrove rivulus can self-assess their physiological tolerance of aquatic hypoxia and make adaptive anticipatory decisions, as the concentrations of oxygen that elicited emergence were ~3-8 fold higher than the concentrations that these fish could tolerate for 20-40 min in our LOE experiments. The mechanism(s) used for self-assessment of hypoxia sensitivity are not known, but previous work has found that the hypoxic emergence response of this species is regulated by oxygen-sensitive neuroepithelial cells (Regan et al., 2011). One possibility is that self-assessment occurs via internally oriented neuroepithelial cells that sense blood P_O2_ (Perry et al., 2009), as impaired respiratory function in air-acclimated fish would presumably cause a faster decrease in blood P_O2_ compared to water-acclimated fish. Understanding the mechanism of self-assessment is an exciting area for future research.

### Establishment of an experimental positive feedback loop

Our habitat choice experiment provides further evidence for positive feedback between gill plasticity and habitat choice. We found that fish acclimated to extreme aquatic hypoxia spent substantial amounts of time out of water (~30% of the experiment, 70-fold more than controls in normoxic water) and were more sensitive to aquatic hypoxia than fish acclimated to normoxia or moderate hypoxia. Fish in the extreme aquatic hypoxia group were also less hypoxia tolerant (i.e. time to LOE was reduced by ~40%) and had reduced functional gill surface area. Notably, these plastic changes are opposite to those typically observed with hypoxia acclimation in fishes (Perry et al., 2009; Richards, 2009). For example, when mangrove rivulus were prevented from leaving water during hypoxia acclimation, respiratory function improved and gill surface area increased (Turko et al., 2012). Overall, our results indicate that when mangrove rivulus are given the choice between living in extremely hypoxic water or on land, the respiratory phenotype is shaped by the terrestrial habitat. Thus, behavioural habitat choice may be a key factor that determines physiological tolerance of extreme environments.

While fish in the extreme aquatic hypoxia treatment spent much more time out of water than fish in the other treatments, as expected, we did not find any change in emergence rates over the course of the 7d experiment. This result is inconsistent with the positive feedback hypothesis, which predicts an increase in emergence over time. Emergence was also surprisingly low in the moderate hypoxia treatment considering that that this oxygen concentration (2.1 kPa) acutely elicited an emergence response in our experiments and is lower than the critical oxygen tension of this species (Turko et al., 2012). Wild mangrove rivulus under similarly hypoxic conditions spent ~90% of their time on land (Turko et al., 2018). One explanation for the low rates of emergence in our experiment was the extensive use of aquatic surface respiration (Chapman and Mckenzie, 2009; Kramer and McClure, 1982). At the levels of aquatic hypoxia we used, mangrove rivulus spent more than 80% of the time displaying this behaviour. We suggest that this extensive use of aquatic surface respiration may be a laboratory artefact, as in the wild ASR exposes fish to predation (Domenici et al., 2007; Kramer, 1983; Riesch et al., 2010) and mangrove rivulus do not appear to frequently use this behaviour in the wild (A. Turko, personal observation). Another possibility is that the low rates of emergence at moderate hypoxia occurred because our experimental conditions were more benign than typical conditions in the field. In addition to hypoxia, wild mangrove rivulus are exposed to intraspecific aggression, high temperatures, hypercarbia, and high levels of hydrogen sulfide; variables all known to cause fish to leave water (Cochrane et al., 2019; Gibson et al., 2015; Robertson et al., 2015; Taylor, 1990). These additional factors may additively increase the overall amount of time rivulus spend out of water. Nonetheless, our results illustrate the interactive effects of habitat choice and physiological plasticity. Even subtle habitat choice, i.e. a shift up in the water column to use ASR, minimized the severity of hypoxia experienced by mangrove rivulus and blunted the plastic responses to gill morphology and respiratory function that occur when fish cannot access the air-water interface.

We noticed a high degree of variation in the tendency of individual fish to leave water in our long-term behaviour experiment, especially in the extreme hypoxia treatment. A similar degree of individual variation in emergence behaviour has been previously reported for this species (Turko et al., 2011). We hypothesise that positive feedback between habitat choice and phenotypic plasticity acts to accentuate individual differences in behaviour and may in part explain the high degree of variation we observed. For example, some individuals may be slightly more likely to leave water as adults (e.g. due to developmental plasticity or epigenetic effects). This behaviour in turn causes an increase in the size of the ILCM and may therefore strengthen the tendency to emerge from water, which could ultimately lead to highly divergent behavioural phenotypes among individuals. Previous work has shown that relatively bold birds choose habitats with higher levels of disturbance, and these habitats may reinforce bold phenotypes (Holtmann et al., 2017). Currently, little is known about how matching habitat choice and variances in individual behaviour affects ecological outcomes, but we think this is an important area for future work.

### Perspectives

A fundamental goal of physiological ecology is understanding the processes that promote phenotype-environment matching (Bolnick and Otto, 2013; Edelaar et al., 2008). At the level of individual animals, the two most important mechanisms are behavioural habitat choice and phenotypic plasticity. These mechanisms are often studied in isolation, and modelling studies have even suggested that selection should favour one or the other depending on the type of environmental variation an animal experiences (Botero et al., 2015; Edelaar et al., 2017; Scheiner, 2016). However, our results combined with other empirical studies (Boyle and Start, 2020; Lowe and Addis, 2019) indicate that plasticity and matching habitat choice interact and even positively feed back on one another. Understanding these interactions can be important for interpreting patterns of phenotypic variation and predicting how animals will respond to changing environments. For example, our data showing that hypoxia-acclimated mangrove rivulus have relatively low gill surface area and hypoxia tolerance cannot be explained without considering both plasticity and habitat choice. Understanding the conditions that give rise to habitat choice-plasticity feedback interactions versus habitat choice or plasticity alone is an important area for future research.

Positive feedback moves systems away from equilibrium and as a result this feedback been hypothesized to facilitate major evolutionary transitions (Crespi, 2004; Hendry, 2016) such as the invasion of land. For example, on evolutionary timescales, phenotypic change coupled with matching habitat choice is thought to promote local adaptation and potentially speciation (Camacho et al., 2020; Jacob et al., 2017; Muñoz and Losos, 2018; Nicolaus and Edelaar, 2018). Our results indicate that analogous feedback between plasticity and habitat choice can cause phenotypic divergence within the lifetime of individuals. Given that expressions of plasticity can cause evolutionary change through genetic assimilation (Crispo, 2007; Pigliucci et al., 2006; Schneider and Meyer, 2017), we speculate that positive feedback between habitat choice and plasticity may be an important first step in generating similar evolutionary feedback between these processes. For example, populations of mangrove rivulus that inhabit moderately versus severely hypoxic habitats should differ in emergence rates and thus respiratory phenotype. If these environmental differences persist, evolutionary processes (e.g. genetic assimilation and/or the altered selective environment) may reinforce the phenotypic divergence between populations. Over time, this mechanism may therefore promote a major ecological transition by producing terrestrially adapted fishes that have poor aquatic performance due to respiratory trade-offs.

## Acknowledgements

We thank Matt Regan, Giulia Rossi, Louise Tunnah, and Paige Cochrane for helpful discussions, and Matt Cornish, Mike Davies, Abiran Sritharan for animal care. This work was supported by the National Sciences and Engineering Research Council of Canada (NSERC) graduate scholarship to AJT and an NSERC Discovery Grants to PAW.

## Notes

### Competing Interest Statement

The authors have declared no competing interest.

